# Torsional Force by Helical Pericytes Regulates Blood Flow in Downstream Capillaries

**DOI:** 10.64898/2025.12.01.691715

**Authors:** Gülce Küreli, Nevin Belder, Meftun Doğa Başaran, Mahmoud Haddara, Luis Alarcon-Martinez, Şefik Evren Erdener, Turgay Dalkara

## Abstract

This study investigates the contractile properties of downstream capillaries, which are traditionally regarded as passive conduits, and addresses ongoing debates surrounding blood flow regulation. It is widely accepted that these capillaries passively dilate in response to increased blood flow in upstream microvessels, and that the helical pericytes located on them lack contractile capability, largely due to the absence of detectable α-SMA expression.

Challenging this prevailing view, we demonstrate that downstream capillary pericytes do express both α-SMA and Myosin11, as shown using in situ hybridization on whole-mount intact retinas—unlike prior studies that relied on dissociated cells. Furthermore, Förster resonance energy transfer (FRET) analysis reveals that α-SMA and Myosin11 are in sufficiently close proximity to permit actomyosin bridge cycling, a process essential for contraction. We also show that pericyte contraction can be inhibited by disrupting this molecular interaction.

Distinct from the nodal constrictions caused by circular pericyte processes in upstream microvessels, we identify torsional contractions in the downstream capillaries in the retina of living mice using two-photon laser scanning microscopy (TPLSM), which regulates blood flow. These contractions provide direct evidence that downstream capillaries actively contribute to blood flow regulation. Notably, such contractions were overlooked in previous TPLSM studies that monitored only luminal diameter, unless the specialized analytical techniques we employed were applied. Our 3D modeling confirms that these torsional contractions correlate with the helical morphology of pericytes in downstream capillaries—a structure previously thought incapable of producing significant constrictive force. In conclusion, our findings provide direct evidence that downstream pericytes play an active role in regulating blood flow. They highlight a previously unrecognized mechanism—torsional contraction—that aligns with the helical structure of these pericytes and contributes to flow regulation in small-caliber capillaries located nearest to regions of high oxygen demand.

**TEASER:** Downstream Capillary Pericytes Express α-SMA and Regulate Flow via Torsional Contraction

## INTRODUCTION

Several recent *in vivo* studies have presented compelling evidence supporting the contractility of pericytes located at the upstream segments of the cerebral and retinal microcirculation (*1–4*). These findings find reinforcement in the histological demonstration of the contractile protein α-SMA in these pericytes (*5–8*). However, the debate over the contractility and α-SMA expression of downstream capillary pericytes persists (*1, 8–10*). The slender, strand-like processes of downstream midcapillary pericytes are believed to lack the proper form for effective contraction. Moreover, the controversy is compounded by the absence or minimal expression of α-SMA mRNA in dissociated vascular mural cells (*1, 9–13*). It is generally accepted that the initial capillary branches off the arteriole can regulate blood flow to the downstream capillary bed in response to electrochemical signals conducted from the capillary bed closest to active neurons (*1, 2, 4, 14*). Consecutive bifurcations of upstream capillaries with contractile ability can serve redirecting the flow to the active focus (*1*). Notably, downstream capillaries form loops with common edges. Computational analyses have demonstrated the universality of this loop structure across species and tissues (*15*). The size and shape of the loop ensure the necessary oxygen gradient to the surrounding tissue from downstream capillaries having relatively lower pO_2_ compared to upstream ones. While past assumptions regarded the changes in red blood cell (RBC) flux within the loops as passive (*10, 14*), emerging evidence now suggests that subtle but active diameter changes may also contribute to precise adjustment of blood flow to meet the ever-changing focal demand while simultaneously sending dilatory signals upstream (*2, 3, 14, 16–18*). For instance, a recent study documented that almost half of the capillaries connected via inter-pericyte tunneling nanotubes constricted while others dilated in response to light stimulation of the retina (*5*). Inter-pericyte tunneling tubes appear to mediate this opposing response by connecting pericytes located in distal capillaries.

In a series of experiments conducted in our laboratory, it was revealed that α-SMA in downstream small pericytes undergoes rapid depolymerization during tissue fixation, thereby eluding detection through histological methods (*8, 19, 20*). As a well-established phenomenon in the extensive actin literature, monomeric (globular) actin (G-actin) is recognized as a stable protein with a prolonged half-life (slow turnover). However, the pool of actin filaments (F-actin) is dynamically replenished through the de/repolymerization of actin monomers, a process that relies on ATP and is regulated by local cellular needs (*21*). Consequently, transgenic mice expressing fluorescent reporter proteins under the Acta2 promoter may have underestimated the presence of α-SMA protein in small downstream pericytes (*1, 22, 23*). The detection of α-SMA protein is further complicated in the immunostaining of thick tissue slabs required to visualize all capillary branch orders due to rapid depolymerization during tissue processing, particularly with slow acting aldehyde fixatives such as PFA. Indeed, the use of rapid tissue fixation methods or depolymerization inhibitors revealed that even high-order downstream pericytes in the retina including those in intermediate and deep layers express α-SMA, albeit in small amounts (*5–7, 24*). This conforms to the ubiquitous expression of Myosin11 in pericytes located on both upstream and downstream microvessels (*1, 23, 25*)

To elucidate whether downstream pericytes on high branch order capillaries exhibit functional contractility, our study focused on three key aspects: co-expression of α-SMA and Myosin11, functional actomyosin cross-bridge cycling, and the organization of contractile fibers (circular vs. strand-like) to discern different contraction phenotypes and assess their potential impact on blood flow regulation. The inherent challenges posed by the small size of downstream capillaries, brain and retina pulsations, and optical resolution limitations have precluded a direct *in situ* investigation of these parameters using two-photon microscopy. Therefore, by leveraging whole mount retina, intravitreal drug administration in vivo, and employing snap tissue fixation techniques, we successfully demonstrated the co-expression of α-SMA and Myosin11 and their mRNAs in both downstream and upstream pericytes. Furthermore, utilizing Förster resonance energy transfer (FRET) analysis, we revealed that the interaction sites of α-SMA and Myosin11 are sufficiently close to enable actomyosin bridge cycling. Inhibition of actomyosin bridge cycling by blebbistatin, an established inhibitor of actomyosin coupling (*26*), effectively prevented contractility. Using 3D models of capillaries, we unveiled for the first time that axial and oblique α-SMA fiber bundle orientations in downstream pericytes resulted in twisting contractions of the vessel wall unlike the nodal constrictions induced by the circular pericyte processes in upstream vessels. Notably, we confirmed the torsional contractions of downstream capillaries in the retina of living mice with two-photon laser scanning microscopy (TPLSM), providing direct evidence of their functional role in blood flow regulation. These comprehensive investigations collectively shed light on the contractile nature of downstream pericytes, revealing their distinctive torsional forces that align with their helical structure.

## RESULTS

### Up and downstream capillary pericytes express Acta2 and Myh11 mRNA

We employed the RNAscope *in situ* hybridization technique to evaluate the expression of Acta2 (gene coding for α-SMA) and Myh11 (gene coding for Myosin11) in both upstream and downstream pericytes along the microvascular tree in the mouse retina. The selection of the Myosin11 isoform of myosin II was based on pericyte single-cell transcriptomics data ((*27*); http://mousebrain.org/) and supported by a recent report (*1*). The use of *in situ* detection in intact tissue was crucial due to the variable morphological and functional phenotype exhibited by pericytes along the microvasculature, making it challenging to discern location-specific transcriptional differences through scRNAseq (*28, 29*). This difficulty arises from technical limitations in dissociating pericytes from the vessel wall and the absence of definitive location-specific pericyte markers (Hartmann et al., 2015). The analysis revealed abundant transcription of Acta2 and Myh11 in the upstream branches of the retinal capillary network (0-2^nd^ branch orders, when the radiating arteriole taken as 0). Transcripts were localized in the peri-nuclear cytoplasm as well as in processes (Figure 1). In more than 2/3 of the downstream branches covered by capillary pericytes, both transcripts were detectable predominantly in the peri-nuclear cytoplasm. The decline in transcript numbers toward downstream branches (3–5) was statistically significant for both genes within the peri-nuclear cytoplasm as well as proximal and distal pericyte processes (Jonckheere-Terpstra Test, p < 0.001). The transcription levels of Acta2 and Myh11 genes exhibited a high correlation (r_s_ (84)= 0.855, p < 0.001)(Figure 1D). We were able to detect both transcripts concurrently in several midcapillary pericytes in the intermediate layer as well (Fig 1C_III_). However, the distribution of positive pericytes was not homogeneous and was scarce, given that 60% of pericytes in this layer are α-SMA-positive (see Figure 3D of Alarcon-Martinez et al.), increasing the possibility of false-negative results due to uneven probe penetration. More aggressive treatment conditions to enhance probe penetration were not tolerated by the thin retinal tissue. Altogether our findings indicate co-expression of α-SMA and Myosin11 in pericytes on downstream capillaries, mirroring the expression patterns observed in upstream pericytes. The observed decline in transcript levels in downstream branches aligns with our previous immunohistochemical findings illustrating α-SMA and Myosin11 protein expression in the retina (*8, 23*).

**Figure 1:**
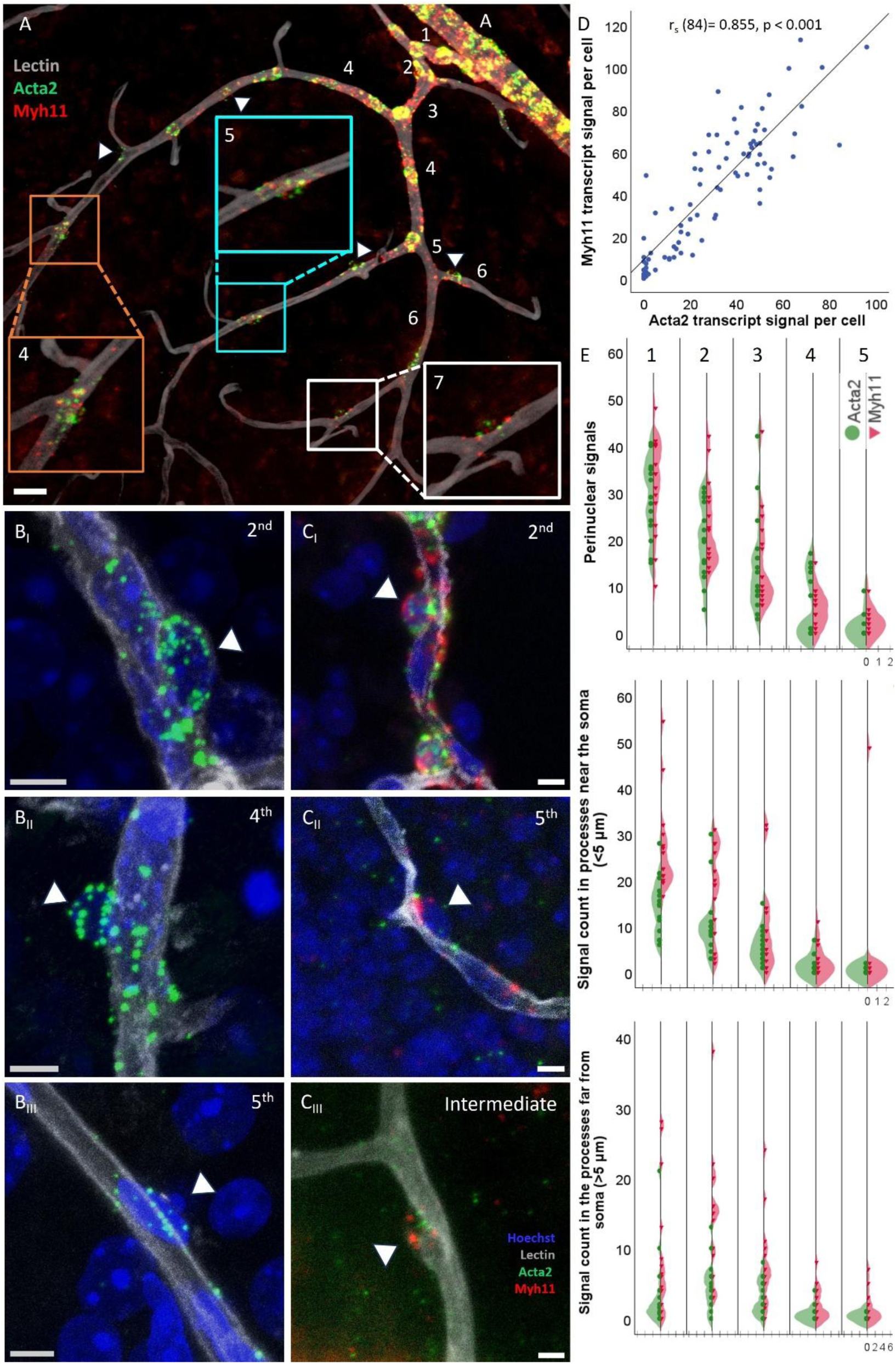
Upstream and downstream capillary pericytes express Acta2 and Myh11 mRNA A) Pericytes along all retinal microvascular orders are labeled for Acta2 (green) and Myh11 (red) transcripts using in situ hybridization. Scalebar: 20 um. Acta2 and Myh11 transcripts are abundantly expressed in the first four capillary branches (counting from the radiating arteriole, PA) of the superficial retinal layer. Vessel walls are delineated with lectin staining of the basement membrane (gray). Transcripts localize in the peri-nuclear cytoplasm in soma as well as in processes. Colocalization of both transcripts produces the yellow color in this merged image of red and green channels, particularly in the pericyte soma. Transcript abundance diminishes toward downstream branches but remains detectable down to the 7^th^ order. Insets are magnified x2 to better illustrate the transcripts in downstream pericytes. Representative maximum intensity projection images from 2^nd^-5^th^ order branches depict Acta2 (green) in images B_I-III_. Panels in column C illustrate both Acta2 (green) and Myh11 (red) and are standard deviation projections of z-stacks for the 2^nd^ (C_I_) and 5^th^ (C_II_) orders, while the intermediate layer image (C_III_) is a maximum intensity projection. Pericyte nuclei (blue, identified with Hoechst staining) protruding out of the vessel wall are marked with arrowheads in B and C. The last panel in C depicts a capillary from the intermediate layer, expressing both transcripts which help identify pericyte soma. Scalebars: 5 um. D) The transcript levels of Acta2 and Myh11 genes exhibit a high correlation [r_s_ (84)= 0.855, p < 0.001]. E column: Graphs illustrate the number of Acta2 (green circles) and Myh11 (red triangles) transcripts within the peri-nuclear cytoplasm as well as proximal and distal pericyte processes. Transcript signals are counted as fluorescent dots within the compartments in relation to the distance from pericyte somas. X-axes show the number of cells, with one cell increments between each tick, having the certain number of signals displayed on the y-axes. The midline between the two transcript signals shows the axis origin (zero) for each vascular order, the numbers increasing as positive integers in both directions. The decline in transcript numbers toward downstream branches was statistically significant for both transcripts (Jonckheere-Terpstra Test, p < 0.001).

### α-SMA and Myosin11 proteins are highly colocalized in up and downstream pericytes

Given the shared features between pericytes and upstream vSMCs, including the co-expression of Myh11 and Acta2 mRNA, it is plausible that pericyte contraction is also mediated by actomyosin cross-bridge cycling. We first investigated the co-localization of myosin II with α-SMA in pericytes with immunohistochemistry (IHC). Our findings showed that Myosin11 was consistently expressed in all vascular orders and retinal layers unlike α-SMA immunoreactivity (Figure Suppl. Fig. 1). Where present, both immunoreactivities exhibited a high degree of overlap, strongly suggesting tight co-localization of both proteins (see Figure 2). It’s worth noting again that rapid fixation with methanol was essential to prevent α-SMA depolymerization during tissue processing as reported before (*8*).

**Figure 2:**
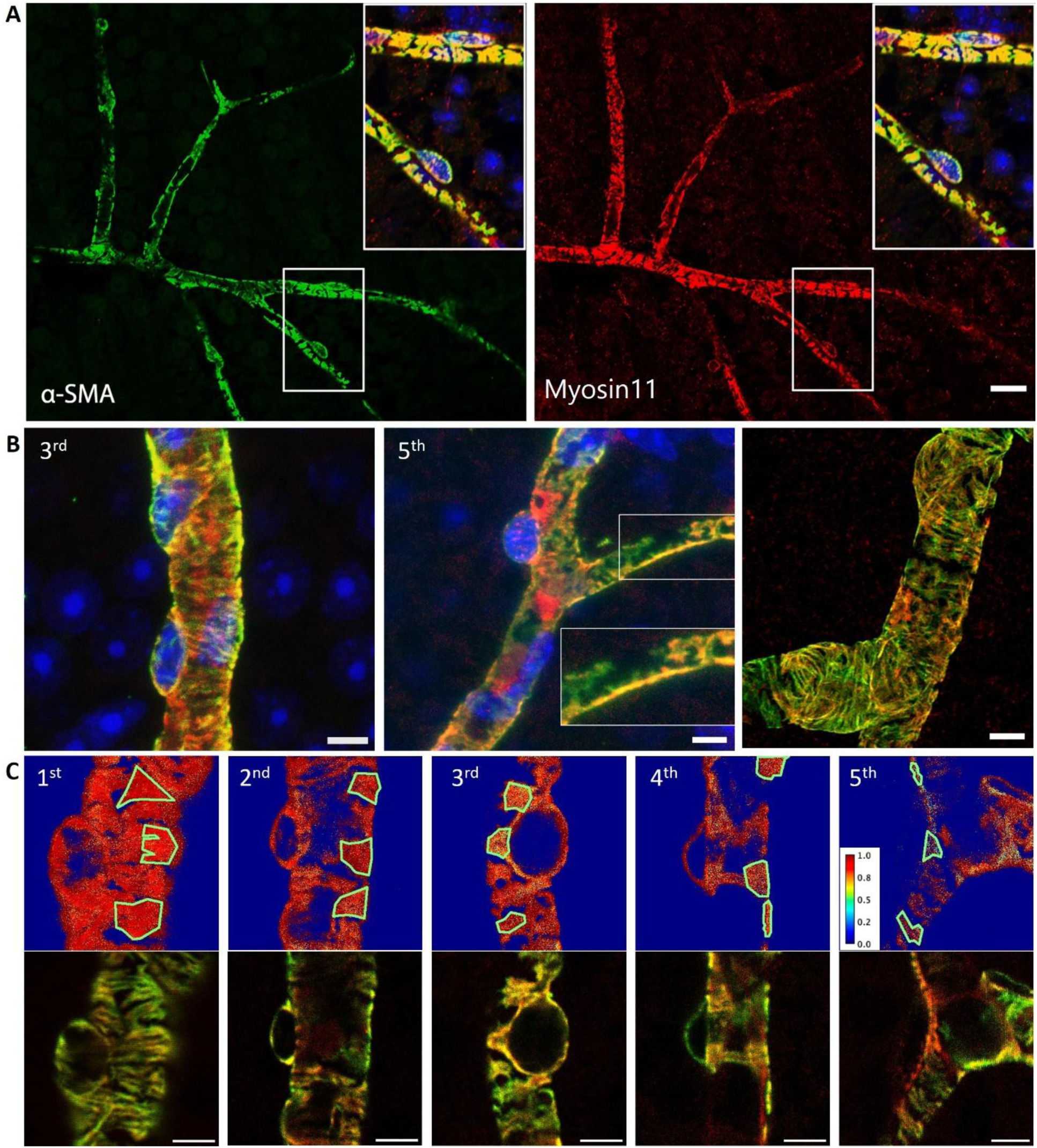
Tight Colocalization of α-SMA and Myosin11 Proteins in Upstream and Downstream Pericytes is confirmed by FRET A) Myh11 (red) and α-SMA (green) immunolabeling exhibit tight colocalization in mesh-type pericytes. Pericytes in the zoomed in insets are located on 3^rd^ order branches. The intense yellow color in merged images signifies the close overlap between the two proteins. Zoomed insets are magnified 2x and display merged images of the green and red channels along with Hoechst nuclear labeling in blue. Scalebar: 20 µm. B) High resolution images provide finer detail on how these two contractile proteins are organized in the form of fibers in individual pericytes to facilitate contraction. The middle panel illustrates the tight colocalization of two contractile proteins in a thin-strand pericyte located on a 5^th^ order capillary. Scalebar: 5 µm. Inset is magnified 1.5x. C) FRET interactions between Cy3 labeled anti-α-SMA and Alexa Fluor-488-nanosecondary antibody labeled anti-Myosin11 antibodies confirm that the interacting sites of two contractile proteins are close enough to allow functional coupling (Warmer colors indicate higher FRET efficiency). Green-outlined regions delineate the ROIs used for FRET efficiency calculation for each pericyte, as detailed in the Extended Data section. The upper row displays the FRET efficiency maps between the two fluoroprobes in pericytes located on branch orders ranging from 1 to 5. The lower row illustrates the corresponding high-resolution images of double immunostaining. Scalebars: 5 µm.

**Fig 3.**
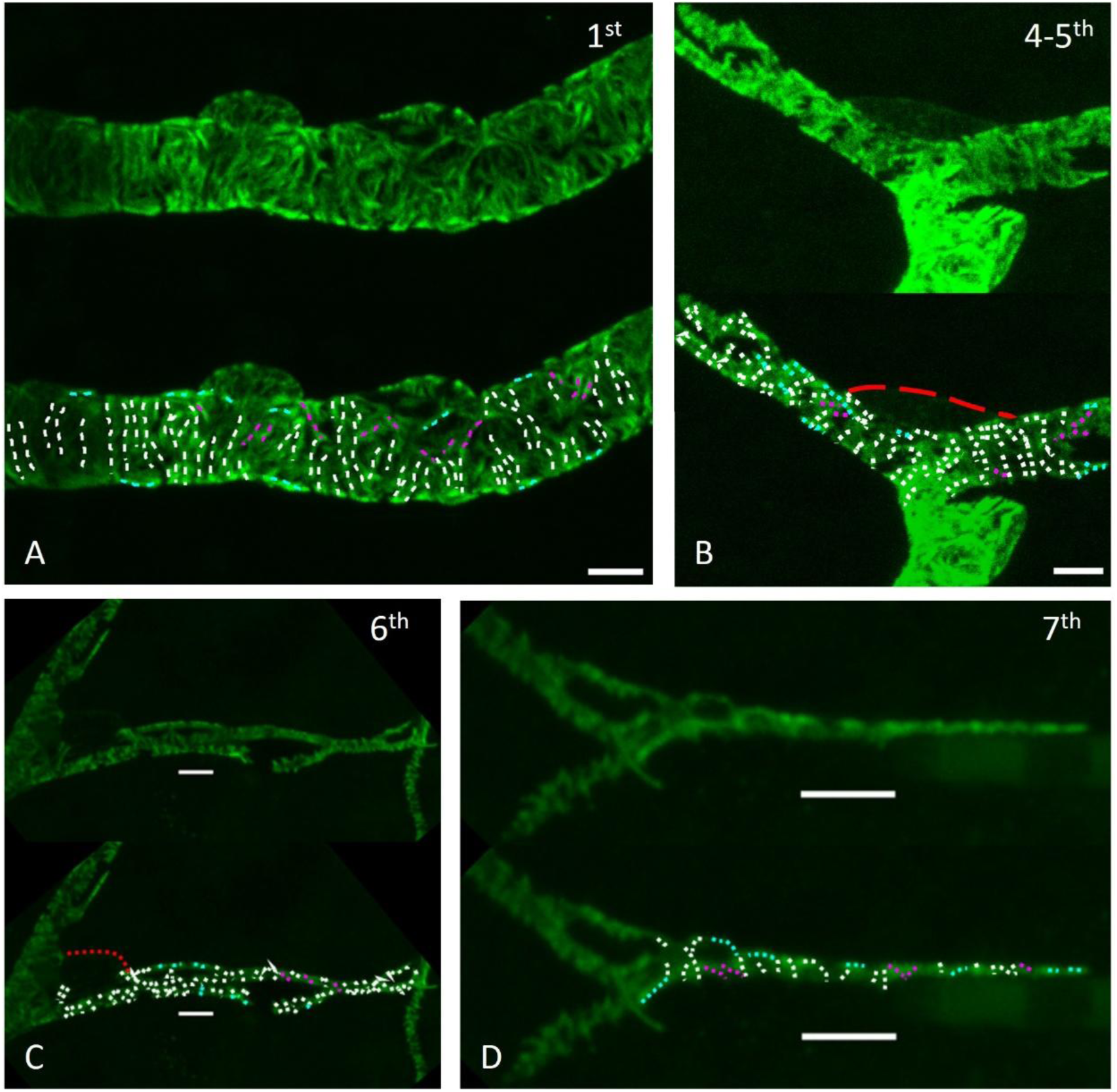
Organization of α-SMA Fiber Bundles in Pericytes for Different Contraction Patterns High-resolution imaging of α-SMA immunoreactivity (green) illustrates the organization and orientation of fiber bundles within the cytoplasm and processes of pericytes across vascular orders. In upstream pericytes (A), the bundles are predominantly circumferential. Towards downstream branches (B), the orientation of the bundles, which are relatively shorter, becomes oblique or parallel to the longitudinal axis of the capillary, especially in the thin-strand processes of 6^th^ and 7^th^ orders (C and D, respectively). Circular bundles are indicated by white dashed lines, oblique bundles by magenta lines, and longitudinal bundles by cyan lines. Pericyte nuclei are outlined in red. Scalebars 5um.

### FRET analysis confirms tight colocalization of α-SMA and Myosin11 in up and downstream pericytes

The close colocalization of α-SMA and Myosin11 immunoreactivity strongly imply a functional interaction between these two contractile proteins. To ensure that their interacting sites are close enough to allow functional coupling, we employed FRET analysis. For this, we used antibodies directed against α-SMA and the head of myosin heavy chain. By spectral unmixing approach (*30*), FRET interactions between Cy3 labeled anti-α-SMA and Alexa Fluor-488-nanosecondary antibody labeled anti-Myosin11 antibodies were evaluated in pericytes (Figure 2C). The results demonstrated a high FRET efficiency above 0.8 in pericytes located on branch orders ranging from 1 to 5 (Suppl. Fig.2), corresponding to a distance of approximately 54 Å (or 5.4 nm) between Alexa488 and Cy3 pairs, based on the Förster critical distance of 67.6 Å for this pair of fluorophores. Pericytes on higher branch order capillaries beyond the 6^th^ could not be examined due to insufficient α-SMA fluorescence signal intensity noted above.

### α-SMA fiber bundles in pericytes are organized to provide different contraction patterns

High-resolution imaging of α-SMA labeled pericytes unveiled the intricate organization and orientation of fiber bundles within both the cytoplasm and processes. The bundles exhibited predominantly circumferential alignment in upstream pericytes, resembling the pattern observed in vSMCs. However, as we progressed towards downstream branches, the orientation of the bundles, which were relatively shorter, became less circumferential. Instead, they ran obliquely or parallel to the longitudinal axis of the capillary (Figure 3). These varying stress fiber orientations prompted us to investigate the constriction patterns in 3D to discern any torsional movement resulting from contraction of the oblique/longitudinal fibers in processes of downstream midcapillary pericytes.

### 3D models of constricted capillaries and α-SMA fibers

We constructed 3D models of capillary lumina, identified through fluorescent-conjugated lectin labeling along with the associated pericytes, marked with α-SMA immunostaining. To induce pericyte contraction, we administered NA into the vitreous body, avoiding systemic effects and trauma to the retina. Although NA’s site of action (e.g., pericytes, astrocytes or endothelia) is not characterized for retinal microvessels, we opted for NA based on our previous observations that intravitreal administration of 1 μL of 100 μM NA significantly reduced capillary luminal diameter, as assessed by lectin labeling (*19*). Two distinct contraction patterns were observed: one featured a marked decrease in luminal diameter from the periphery of the pericyte towards the soma (node-like) (Figure 4A). In this pattern, the capillary wall moved towards the center of the lumen, with maximal constriction occurring under the soma. The second pattern involved a subtle displacement (indentation) of the soma into the lumen, accompanied by a gradual but diffuse reduction in diameter along the primary pericyte process (tide-like) (Figure 4A). These *ex vivo* observations of retinal capillaries, perfusion-fixed under *in vivo* conditions after intravitreal NA application, closely resemble descriptions of pericyte contraction induced by focal electrical stimulation along capillary segments *ex vivo* in whole mount retina preparations (*31*). Node-like contractions were exclusively identified on proximal segments covered with mesh-type pericytes, while tide-like contractions were observed on downstream segments covered with midcapillary pericytes.

**Fig 4.**
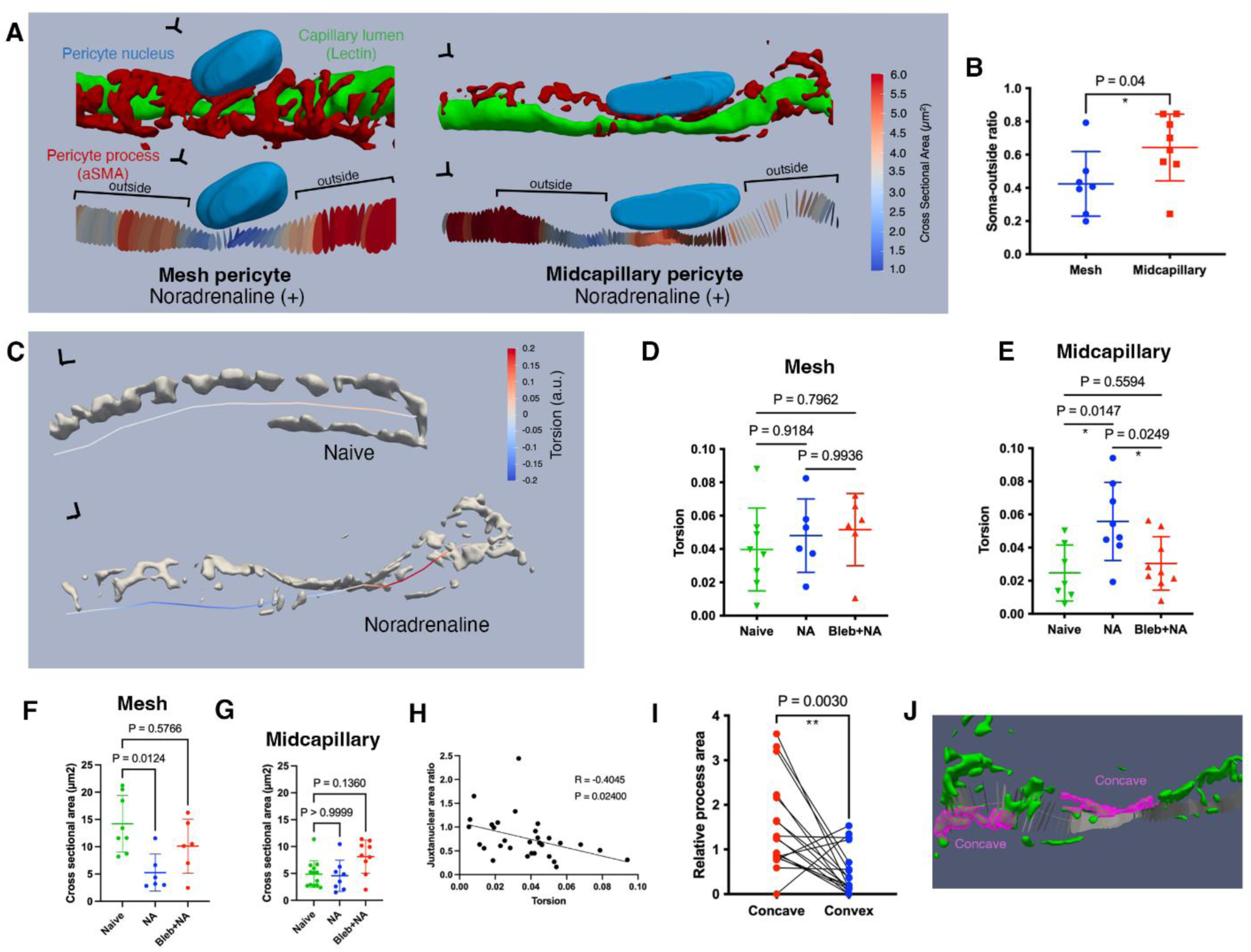
3D analysis of ex vivo images reveals a torsional component in pericyte contraction (A) Comparison of ex vivo 3D models and cross-sectional area profiles of two representative capillaries with mesh type and midcapillary pericytes, from noradrenaline-treated retinas. Pericyte processes, lumen and nuclei were segmented and reconstructed from α-SMA (red), FITC-Conjugated-Lectin (green) and Hoechst (blue) fluorescent signals, respectively. Mesh pericytes with circumferential processes cause a sharp decrease underneath pericyte soma, compared to regions outside the soma, while midcapillary pericytes result in a subtle but diffuse (tide-like) decrease in lumen, including the areas outside the soma. Scale axes: 1 µm. (B) Quantitative data show that cross-sectional diameter ratio between the soma and outside-soma regions is significantly lower in noradrenaline-treated capillaries with mesh type pericytes compared to those with midcapillary pericytes (Mann Whitney U Test, P=0.04). (C) Centerline torsional profiles of representative capillaries with midcapillary pericytes from naive and noradrenaline-treated retinas, disclose both clockwise (positive, red) and counterclockwise (negative, blue) torsional movements. Scale axes: 1µm. (D,E) While median torsion in each segment does not significantly differ between noradrenaline, noradrenaline+blebbistatin (Bleb) treated, and naive groups in upstream capillaries. However, in midcapillary pericytes, noradrenaline induces a significant increase in torsion, which is prevented by actomyosin coupling inhibitor blebbistatin (Kruskal Wallis Test followed by Dunn multiple comparisons, P<0.05). (F,G) Conversely, noradrenaline causes a significant decrease in median cross-sectional areas of upstream capillaries, which is also prevented with blebbistatin, whereas median cross-sectional areas of downstream capillaries harboring midcapillary pericytes show no significant change (Kruskal Wallis Test, followed by Dunn multiple comparisons). (H) Increase in median torsion in each capillary segment shows a significant negative correlation with juxtanuclear area ratio, that is the ratio of the median cross-sectional area next to but not underneath the nuclei to the the median cross-sectional area closest to the origin of the segment and distant to the pericyte nuclei (Pearson Correlation, R=-0.4045, P=0.024). (I-J) Torsional bends are driven by reciprocal organization of processes of midcapillary pericytes. The concave sides of torsional bends of the capillaries have significantly higher area of pericyte coverage compared to the convex sides of the bends (Paired t-test, P=0.0030).

These two distinct constriction patterns were quantified by comparing the ratio of the minimum cross-sectional area under the soma to the median cross-sectional area outside but within a two-nucleus distance of the soma. This ratio indicated how sharply the luminal cross-sectional area diminished towards the pericyte soma from the periphery (Figure 4B). In accordance with the visual observations, a significantly lower ratio was detected for mesh pericytes compared to midcapillary pericytes in NA-treated retina (Figure 4B).

### Midcapillary pericytes exert torsional forces via actomyosin crossbridge cycling

Leveraging the advantages of 3D capillary reconstructions, we computed the centerline torsions of capillaries covered by mesh and midcapillary pericytes in both the naive and NA-treated retina (Figure 4C). For capillaries covered by mesh pericytes, torsion measurements showed no significant difference between the naive and NA groups (Figure 4D). However, a notable increase in torsion was observed in capillary segments covered by midcapillary pericytes on the 5^th^ and 6^th^ order capillaries from the NA-treated retina compared to the naive group (Figure 4C, E). Importantly, this increase was prevented by intravitreally introducing blebbistatin before NA, suggesting that the torsional contraction was mediated by actomyosin coupling (Figure 4E). Despite the evident torsional contraction, the luminal cross-sectional area of the capillaries covered by midcapillary pericytes was not appreciably reduced after NA, in contrast to upstream capillaries covered by mesh pericytes, which exhibited blebbistatin-sensitive circumferential constriction (Figure 4F,G). In a previous study (Kureli et al., 2020), we found that the juxtanuclear diameter ratios of capillaries were modulated by both noradrenaline and blebbistatin. Consistent with this, juxtanuclear cross-sectional area ratios in the 3D capillary models—a comparable metric—showed a significant negative correlation with torsion (Figure 4H). This ratio also trended lower in NA-treated retinas compared to naïve retinas, although the difference did not reach statistical significance, possibly because the exact junctional origins of the modeled segments were not within the imaging field of view, due to the magnification and resolution required for precise 3D modeling. Altogether these observations suggest that both downstream and upstream pericytes contract by actomyosin coupling but downstream capillaries with midcapillary pericytes exhibit torsional as well as tide-like contraction patterns, conforming with the stress fiber orientations described above.

Subsequently, we investigated the relationship between pericyte process organization and capillary torsional bends in mid-capillaries. We compared the pericyte process coverage on the convex and concave parts of the segments with respect to the torsional bends detected. We found that the concave sides exhibited significantly higher pericyte process coverage compared to the convex sides (Figure 4I,J). This finding suggests that capillary torsion is driven by the asymmetrical arrangement of pericyte processes in the helical strand surrounding that particular segment.

### Two-photon imaging confirms midcapillary torsional changes that regulate blood flow in living mice

Next, we corroborated this ex vivo observation with two-photon microscopy imaging of the vascular plexuses of the retina in living mice (Fig. 5A). Importantly, we selectively visualized the intermediate and deep vascular plexus as they are exclusively formed by high branch order vessels with midcapillary pericytes. We non-invasively imaged through the sclera of the intact eyeball after intraperitoneal injection of fluorescein to label the plasma (Fig. 5B-C). Computing centerline torsion from high-resolution images of distal capillaries provided the first evidence, to our knowledge, of the torsional contractions of downstream capillaries in living mice (n=4) (Fig. 5D, E). 3D volumetric reconstructions of the same capillary from repeated two-photon Z-stack acquisitions over minutes documented spontaneous and transient torsional changes in capillary geometry at the same point. Notably, the average luminal diameters of these capillaries, as measured from two-dimensional maximum intensity projection images, showed only modest changes at the limits of the light microscopy resolution. For example, Point X indicated in Fig. 5D had a diameter of 4.38, 4.35 and 4.42 um at the 0, 5, and 9-minute timepoints, respectively, as measured automatically using VasoMetrics toolbox (*32*) (Fig. 5D).

**Fig. 5.**
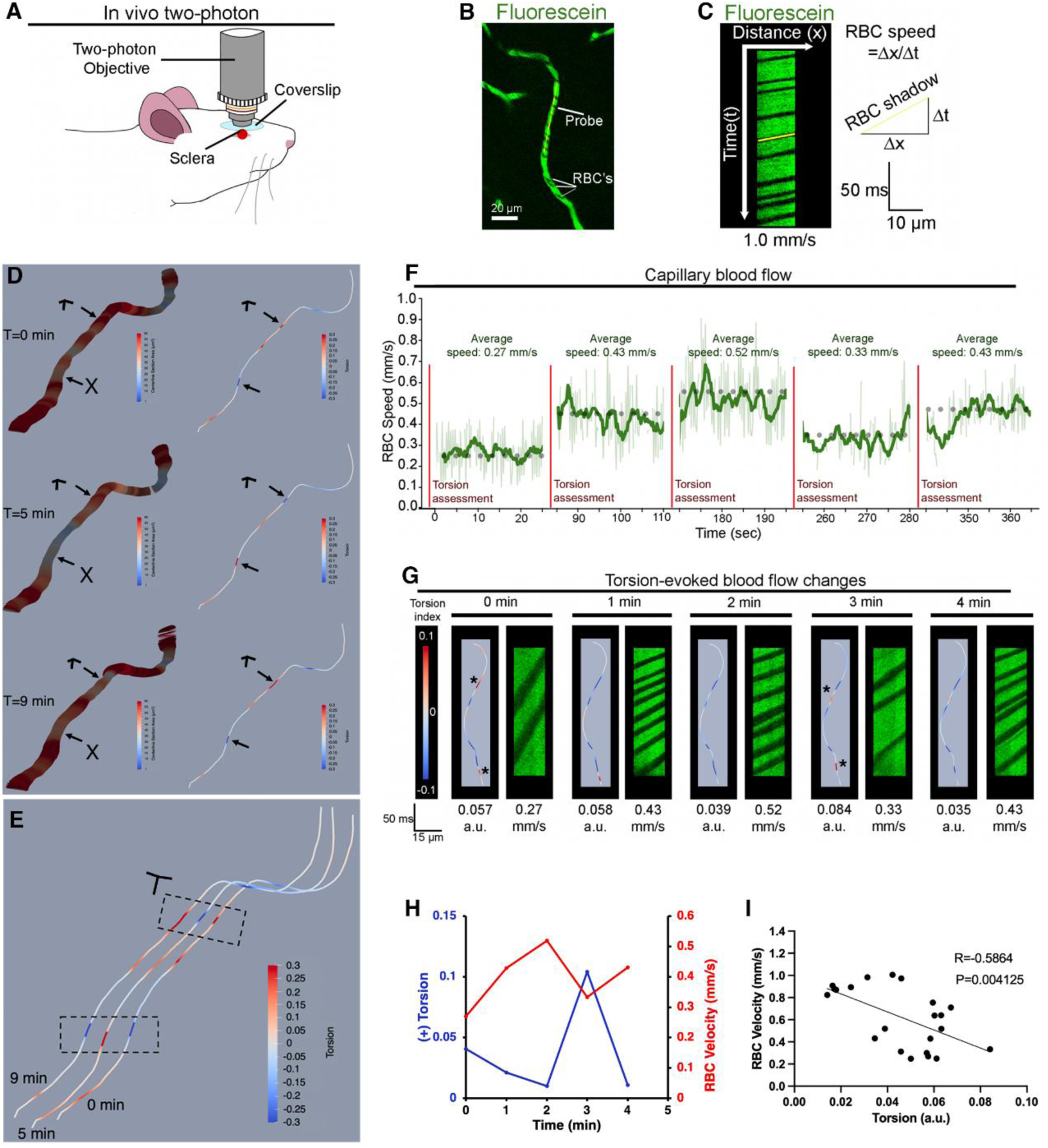
3D analysis of two-photon microscopy images of retinal capillaries confirms torsional modulation and its association with blood flow in vivo (A) *In vivo* set-up to visualize the retina with two-photon laser scanning microscopy (TPLSM). (B) Retinal capillaries visualized with intraperitoneally applied fluorescein, enabling the identification of single red blood cells (RBCs). The yellow probe in the center of the capillary lumen indicates the two-photon longitudinal line scan. (C) Diagram depicting line-scan producing distance-time images, where RBCs appear as oblique shadows whose angle is used to calculate RBC speed in mm/s. (D) 3D volumetric reconstructions of the same capillary from repeated two-photon Z-stack acquisitions over 10 min reveal spontaneous torsional changes in capillary geometry. Cross-sectional area profiles (left column) and centerline torsion profiles (right column) are shown 5 and 9 min after the beginning of data acquisition. Arrows indicate the hotspots for torsional changes at 5 min, which reverse at 9 min. Scale axes: 5 µm. (E) Side-by-side view of the three centerline torsion profiles for enhanced clarity. Scale axes: 5 µm. (F) Kymograph of the capillary depicted in (B) shows changes in RBC speed over 5 minutes across five separate measurements. The time-points of torsional measurements using volume stacks is highlighted with red vertical lines. (G) Illustration of torsion-evoked blood flow changes in distance-time images with corresponding torsional changes in the same capillary shown in F. (H) In the same capillary, average positive torsion changes (blue) show an inverse association with corresponding average RBC velocities (red) plotted over time. (I) Linear regression of average RBC speed and average torsion across all animals and measurements demonstrate a statistically significant negative linear relationship (Pearson correlation, R=-0.59, p<0.01).

This subtle variation may account for why previous two-photon microscopy studies, which focused primarily on capillary diameter changes, did not detect this phenomenon. Next, we investigated whether the centerline torsional changes regulate blood flow under physiological conditions. By interlacing line-scans with repetitive Z-stacks in retinal capillaries, we monitored spontaneous torsional changes alongside red blood cell (RBC) velocities. In 3 out of 5 capillaries observed (n=3 mice) over 5 min, we detected prominent torsional changes. These torsional values were inversely associated with RBC velocity in the same capillary over time (Fig. 5-F-G-H-I). Pooling data from all capillary volumes imaged (total of 26 volumes from 5 capillaries from 3 mice), we found a significant negative correlation between average torsion and RBC velocity (Fig 5I). These findings support the hypothesis that torsional modulation of distal capillaries, along with the subtle, tide-like contraction pattern driven by helical pericytes, plays a role in blood flow regulation.

### Up and downstream pericyte contraction is mediated by actomyosin cross-bridge cycling

To further scrutinize the above view, we compared prevention of NA-induced contraction by blebbistatin in midcapillary pericytes with upstream ensheathing and mesh pericytes. For this, we assessed the luminal diameter (rather than cross-sectional area or torsion elaborated above) to allow comparison of our results with previous published studies. Intravitreal injection of blebbistatin, administered 40 minutes before NA injection, significantly decreased the luminal diameter of capillaries (Suppl. Fig. 3). Due to high variation in vessel sizes across branching orders, for comparative analysis, the magnitude of constriction was normalized by calculating the ratio of the juxtanuclear radius to the radius at the origin of the capillary (referred to as juxtanuclear diameter ratio, JNDR)(*19*). Following blebbistatin pretreatment, JNDR values were higher (i.e. less constricted) compared to values obtained from retinas injected with the vehicle 40 minutes before NA (means: 0.79 vs. 0.76; SDs: 0.20 vs. 0.19, p < 0.001; n=1026 vs. n=1085 vessels, 6 retinas per group). Upon separate analysis of diameter changes based on pericyte types in the superficial layer of the retina, JNDRs were higher in the blebbistatin-treated group compared to the vehicle-treated group for ensheathing (orders 1-2) (means: 0.74 vs. 0.69; SDs: 0.20 vs. 0.17, p = 0.009, n=180 vs. 145) and mesh pericytes (orders 3-4) (means: 0.81 vs. 0.78; SDs: 0.21 vs. 0.19, p = 0.003, n=88 vs. 100). Fifth-and sixth-order capillaries with thin-strand pericytes exhibited less luminal narrowing and higher JNDR (due to tide-like constriction). However, this effect was comparably mitigated by blebbistatin (means: 0.80 vs. 0.78; SDs: 0.18 vs. 18, p=0.06, n=282 vs. 318), to upstream capillaries. When only capillaries showing JNDR smaller than 0.85 (a threshold corresponding to constriction in 4-5^th^ order capillaries according to (*19*) were evaluated, the blebbistatin treated group showed a higher JNDR (p=0.03), showing the effect of this pharmacologic intervention on noradrenaline responsive capillaries These data suggest consistent actomyosin-coupled contraction across all types of capillary pericytes.

## DISCUSSION

We have demonstrated the torsional (twisting) type contractile capability of midcapillary helical pericytes within the retinal microcirculation, mediated by conventional actin-myosin coupling. This was achieved through three-dimensional geometric analysis of in vivo two-photon and ex-vivo 3D images of downstream capillaries along with the detection of mRNAs and proteins associated with contractile elements, specifically α-SMA and Myosin11 in pericytes. Our findings also revealed the intimate localization of interacting sites between these two proteins at FRET distance, which facilitates the coupling necessary for contraction. Accordingly, inhibiting the actomyosin interaction blocked both constrictive and torsional capillary contractility. Furthermore, the orientation of stress fiber bundles in upstream pericytes was circular, while downstream pericytes predominantly displayed an oblique orientation. This suggests that ensheathing/mesh type pericytes contribute to circular nodal constriction of the vessel lumen, whereas midcapillary helical pericytes induce twisting contractions of the vessel wall in addition to subtle diffuse luminal narrowing. These distinct mechanisms may enable the precise regulation of blood flow by adjusting luminal diameter or capillary resistance/tonus, thereby, RBC flux in upstream and downstream capillaries, respectively.

*In vivo* studies consistently report minor dilations in capillaries of the somatosensory cortex, olfactory bulb, and retina during sensory stimulation (*33, 34*). These diameter changes, approximately 1–2% from baseline, challenge the limits of optical resolution in *in vivo* two-photon microscopy. Despite their subtle nature, these dilations have been demonstrated to be physiologically significant, as evidenced by concurrent increases in RBC flux and a reduction in the frequency of intracellular calcium transients in thin-strand pericytes (*14, 35, 36*). Modeling studies have also predicted the physiological relevance of these small diameter changes (*14, 37*). Although small changes in diameter can significantly modulate flow according to Poiseuille’s law, the twisting contraction of downstream pericytes may further influence capillary resistance by altering luminal torsion or modulating capillary wall tension. This functional adaptation aligns with the helical structure of thin-strand midcapillary pericytes and the oblique orientation of stress fibers, as documented in the present study. The helical wrapping of a thin strand around a capillary, coupled with its subtle contraction, is likely energetically advantageous, particularly in downstream capillaries where lower luminal hydrostatic pressure requires less substantial contractile force compared to upstream capillaries with higher intraluminal pressures. The twisting contraction of downstream capillaries poses further challenge to detect with current two-photon microscopy techniques compared to luminal diameter changes, possibly contributing to the difficulty in associating increases in RBC flux and decreases in intracellular calcium transients in thin-strand (helical) pericytes with subtle capillary diameter changes in previous studies (*14, 36*). Our approach of snap-freezing eyeballs allowed us to visualize the retinal vasculature as *in situ*, leveraging three-dimensional high-resolution confocal microscopy, which was also confirmed with 3D torsion analysis of two-photon images obtained from live mice.

The mechanism described in downstream capillaries may serve a crucial role in the retina and possibly in brain due to their unique metabolic demands. Unlike many other tissues, the retina and brain maintain a high resting metabolism, and their temporary functional activation tends to be focal. The presence of resting capillary flow heterogeneity seems essential as it creates the capacity for functional hyperemia while meeting basal metabolic needs (*38*). In essence, the heterogeneity in capillary flow allows high-resistance capillaries with low flow at rest to significantly dilate during functional activation. This dilation increases the flux rate, matching the heightened oxygen demand in the activated region. For instance, during hypercapnia, smaller capillaries exhibit greater dilation and increased RBC flux compared to capillaries with larger baseline diameters (*39, 40*). The noradrenergic innervation from the locus coeruleus has been proposed as a mechanism to maintain high capillary resistance by constriction (*41*). The baseline constriction may allow capillaries to increase their radii in response to vasodilatory mediators during functional hyperemia (*41*). Additionally, focal functional hyperemia may necessitate constriction of neighboring capillaries in adjacent inactive loops fed by the same upstream arteriole-capillary transition zone (ACT), as well as those with common edges with adjacent loops fed by different ACTs (*15*). Recently identified reciprocal diameter changes between distal capillaries might indeed reflect such a regulatory mechanism (*5*). This potential regulation could involve diverting the flow to the loop feeding the active area while constricting adjacent capillaries off the focus. Furthermore, the contractile capacity observed in downstream capillaries may also be required for restoring basal tonus after passive or active distension during functional hyperemia (*3, 14, 23*).

In addition to compelling evidence supporting the contractile nature of upstream pericytes (*1–4*), recent *in vivo* studies have demonstrated the contractile capacity of downstream capillary pericytes as well (*3*). Optogenetic stimulation induced contraction in high-order capillary pericytes (*3*), and their ablation resulted in dilation of the underlying capillary segment (*42*), highlighting the role of pericyte contractility in maintaining capillary tonus. These small cells having thin processes, can exhibit contractility driven by a limited number of α-SMA fibers observed through electron microscopy (*43*). However, detecting these fibers with IHC has been challenging. Notably, measures preventing F-actin depolymerization or promoting F-actin polymerization during tissue fixation revealed a significant increase in α-SMA immunopositive downstream capillaries. This suggests dynamic regulation of a small pool of α-SMA through de/re-polymerization in the 5^th^ and 6^th^ order pericytes (note that due to aligning the numbering of branch orders with the consensus for the brain, 5th and 6th orders correspond to 6^th^ and 7^th^ orders in the cited work on retina) (*8*). While the contribution of α-SMA polymerization to force generation in vSMCs is well-established (*44*), its role in CNS pericytes under *in vivo* conditions is just beginning to be recognized (e.g., by suppressing it with fasudil) (*3*), despite clear evidence from prior cell culture studies (*45, 46*). The above findings from various laboratories suggest that downstream pericytes can utilize depolymerization to facilitate passive dilation during functional hyperemia. Conversely, polymerization may be crucial for restoring basal tonus after transient dilations and may also contribute to the active constriction of non-focal capillaries, as discussed earlier. Supporting the role of de/re-polymerization in these processes, the contraction of downstream pericytes is reportedly slow during optogenetic stimulation, the restoration of basal tonus is sluggish after CO_2_-induced vasodilation, and calcium changes, which are independent of L-and T-type voltage-gated calcium channels, occur on a slow timescale unlike faster actomyosin interaction kinetics (*3, 47*). Transcriptomic studies revealing relatively higher expression of the actin-associated protein transcripts in downstream pericytes compared to upstream arteriolar vascular SMCs when normalized to Acta2, the α-SMA transcript, further reinforce the significance of α-SMA and its de/re-polymerization (*27*)(Supplementary Table 1).

In midcapillary thin strand helical pericytes, we posit that myosin II forms stable thick fibrils, as demonstrated through IHC in all downstream pericytes. Concurrently, the thin smooth muscle actin (SMA) strands exhibit variable lengths, adjusting to the required capillary tonus. This adaptive feature may serve to minimize ATP consumption during basal tonic contraction (tonus) by avoiding unnecessary actomyosin interaction. The system can temporarily enhance contractile force by incorporating additional α-SMA monomers (i.e., polymerization) when an extra force is needed. While the presence of Myosin11 in thin strand pericytes has faced less objection compared to α-SMA, single-cell transcriptomic studies indicate low Myh11 mRNA levels despite clear immunoreactivity in histological studies. Our *in situ* hybridization data unequivocally demonstrate the presence of α-SMA mRNA in downstream pericytes in amounts comparable to Myh11 mRNA. This suggests parallel expression of α-SMA and Myosin11, despite challenges in detecting α-SMA protein by IHC in tissues fixed with PFA, as discussed.

In conclusion, it appears that blood flow is intricately regulated through diverse mechanisms along the arteriovenous tree to ensure adequate supply of oxygen to the surrounding tissue in response to varying demand. Arterioles can efficiently meet the needs of the capillary-free Krogh’s cylinder around them through O_2_ diffusion alone due to sufficiently high O_2_ they have. In the microvascular bed, upstream dilation becomes crucial to accommodate the heightened demand at up and downstream level. The downstream loops not only passively dilate following upstream dilation but also possess the capacity to fine-tune blood flow, perfectly matching nearby demands by adjusting transit times through high branch order capillaries. Pericytes emerge as key players in adapting to these functions, employing variations in process morphology, stress fiber orientation, and actin de/repolymerization, in addition to conventional actomyosin coupling. Their versatile roles underscore the intricate and dynamic nature of blood flow regulation throughout the vascular network, ensuring the precise delivery of oxygen to tissues in response to ever-changing highly focal metabolic requirements.

## MATERIALS AND METHODS

### Animals

A total of 32 male and female *Swiss albino* mice weighing between 25-40 g were housed in plexiglass cages and subjected to 12-hour light-dark cycles. Food and water were provided ad libitum. The number of animals required was reduced by using the contralateral eye of each mouse for vehicle injection. All animal experiments adhered to relevant guidelines and regulations, and experimental procedures on mice were approved by the Hacettepe University Animal Experimentations Local Ethics Board (Registration numbers: 2021/04, 2021/68). All animal procedures and experiments performed in Australia had ethics approval by the St Vincent’s Hospital (Melbourne) Animal Ethics Committee (AEC) and complied with the Prevention of Cruelty to Animals Act and the NHMRC Australian code.

### Intravitreal Administration

Intraperitoneal injection of 1-1.5 g/kg urethane induced anesthesia, depth of which was monitored through toe pinch reflex and vital parameters. Mice were positioned under a Nikon SMZ 745T stereomicroscope, with the head secured by a nosepiece, allowing movement around the longitudinal axis. Intravitreal injections were performed using a 33 G Hamilton Neuros Syringe (1701 RN). The needle, leaving 1.5 mm outside the protective sleeve, was inserted through the ora serrata to avoid lens and retina damage. Post-vehicle injection, the syringe underwent thorough cleaning with distilled water. Capillary constriction was induced by intravitreal delivery of 1 μL of 100 μM Noradrenaline (Cardenor, Norepinephrine tartrate, Defarma). Considering the mice’s intraocular fluid volume of about 4 μL (*48*), a dilution factor of 5 was estimated to yield an approximately 20 μM intravitreal concentration. Diluted solutions were stored at +4°C, for a maximum of 2 months and protected from light. The contralateral eye received an equal volume of the vehicle (5% dextrose). Blebbistatin (Abcam), a cell-permeable and reversible inhibitor of myosin II ATPase activity (*49*), in powder form was reconstituted in DMSO, stored at-20°C, and diluted to a 250 μM concentration in saline before use. The reconstituted Blebbistatin was equilibrated to room temperature for an hour, and 1 μL of the 250 μM solution (yielding approximately 50 μM final concentration) was delivered intravitreally 40 minutes prior to NA application in relevant groups. Contralateral eyes were injected with the same volume of saline before NA injection as a vehicle. The routine forceps-assisted enucleation technique, prone to increased intraocular pressure leading to retinal detachment and vitreous body escape, was avoided. Removal and immersion into fixative promptly performed within two minutes due to the short half-life of NA (*50*). Mice were euthanized by cervical dislocation after eyeball extraction.

### Whole Mount Retina Preparation

The whole-mount retina technique was preferred as it provides an ideal platform to assess pericytes across all branching orders, as the planar organization of the superficial vascular layer allows for the visualization of arterioles, capillaries, and venules on a single plane, in addition to imaging the capillaries diving into the intermediate and deep plexuses (*51*). Enucleated eyeballs were fixed by immersion in 100% ice-cold methanol. They were kept at-20°C for one hour and then transferred to phosphate-buffered saline (PBS, Sigma) at +4°C until retina extraction. Fixed eyeballs were transferred to a petri dish containing PBS at room temperature for retina extraction. Under a stereomicroscope, skin, appendages, lacrimal gland, and extraocular muscles surrounding the eyeballs were removed with fine scissors. A minimal piece of surrounding tissue was intentionally left for handling with forceps. A hole was created at the corneal margin with a 26 G needle. Using Vannas Spring Scissors (FST), a circular cut along the *ora serrata* was made to remove the cornea, followed by the removal of lens and vitreous body with a fine forceps. Four radial cuts were made with a fine spring scissor to flatten out the retina. After flattening, the retina was detached from the underlying sclerochoroidal tissue with a fine forceps and gentle handling. The separated retina was transferred to a round-bottom centrifuge tube containing 200 μL PBS, for immediate use or storage at +4°C for 3-5 days.

### RNA Fluorescence In Situ Hybridization (RNA FISH)

For RNA *in situ* hybridization, enucleated eyeballs were fixed in 4% paraformaldehyde (PFA) at +4°C for 2 to 3 hours, washed with 2X PBS (0.1% DEPC treated) at +4°C for 6 minutes, and the whole mount retina preparation was carried out as described above. To visualize Acta2 and Myh11 expression in pericytes along the microvascular tree in the mouse retina, RNA fluorescence *in situ* hybridization (RNA FISH) was performed using RNAscope Multiplex Fluorescent Reagent v2 kits (Advanced Cell Diagnostics, 323100). The manufacturer’s protocol was followed with slight modifications. Whole mount retinas, prepared from enucleated eyes, were fixated overnight in 4% PFA solution at +4°C. The next day, retinas were washed with 1X PBST (1X PBS + 0.1% Tween 20) for 10 minutes and then consecutively incubated in 25%, 50%, 75%, and 100% methanol solutions for 5 minutes each at room temperature. After discarding the 100% methanol solution, retinas were washed three times (3 minutes each) with PBST.

The retinas were hybridized with proprietary oligo probes for mouse Acta-2 (Advanced Cell Diagnostics, 319531) and Myh11 (Advanced Cell Diagnostics, 316101-C2), following the provided protocol. Positive controls (Probe-NHP POLR2A, PPIB, and UBC; Advanced Cell Diagnostics, 320901) and a negative control (Probe DapB; Advanced Cell Diagnostics, 320871) were used to assess RNA quality. After hybridization, signals were amplified and developed with OpalTM 570 or 650 fluorophores (1:750 in RNAscope® Multiplex TSA buffer) (Akoya Biosciences, Marlborough, MA, USA) for their respective channels. The wash buffer was discarded after the final wash cycle, and RNA-hybridized retinas were labeled with fluorescein lectin (1:200 in PBS) (Vector Labs, FL-1171-1) before being mounted on glass slides with mounting medium containing Hoechst 33258.

RNA FISH signal counting for Acta2 and Myh11 was performed according to kit manufacturer instructions. Signals were counted in their respective channels in Fiji. The evaluation was conducted in three different pericyte cellular compartments, namely soma, proximal processes (<5 µm distance to soma), and distal processes (>5 µm distance to soma); and per vascular order.

### Fluorescent Labeling of Vessels

Whole mount retinas underwent washing and permeabilization with 0.1% PBS-Triton-X 100 (Merck) three times for 5 minutes each. Fluorescein lectin (Vector Labs, FL-1171-1), diluted in PBS at a concentration of 1:200, was applied to the tissue for 8 hours or overnight at +4°C. After washing with PBS three times for 5 minutes, retinas were mounted on glass slides with mounting medium containing Hoechst 33258. Spacers, created from intentionally broken coverslips, were used to prevent the squeezing of vascular lumina in retinal preparations (*52*).

### Immunofluorescent Labeling

Whole mount retinas underwent a triple wash and permeabilization process with 0.1% PBS-Triton-X 100 (Merck), each step lasting 5 minutes. Background staining was blocked by employing 10% Normal Goat Serum in PBS. The primary antibody against Myosin11 (Abcam, ab82541) was diluted in blocking solution to a concentration of 1:400 and incubated at room temperature for 4 hours. After washing with 0.1% PBS-Triton X (3 times, 5 minutes each), Cy2 or Alexa Fluor-555-conjugated goat-anti-rabbit secondary antibody (Jackson Immunoresearch), diluted in blocking solution to a concentration of 1:200, was applied for 1 hour at room temperature. Retinas were then washed with 0.1% PBS-Triton X three times, 5 minutes each.

For double labeling with α-SMA, two distinct primary antibodies were used considering the fluorescent emission spectrum of the fluorophore used to label Myosin11. When Cy2 was utilized as the fluorophore for Myosin11, α-SMA primary antibody conjugated to Cy3 (Sigma, C6198) was used. It was diluted in blocking solution to a concentration of 1:300 and incubated overnight at +4°C. When Cy3 was employed as the fluorophore for Myh11, 1 μL of an unconjugated primary antibody against α-SMA (Sigma, A5228) was incubated with 1.25 μL of Alexa Fluor 488-conjugated goat-anti-mouse Fab fragment (Jackson Immunoresearch) in 10 μL PBS for 1 hour at room temperature. After the addition of 288 μL of 10% Normal Mouse Serum in PBS to achieve a 1:300 antibody concentration, the mixture was incubated at room temperature for an additional hour. Following the removal of the secondary antibody used for Myosin11 labeling, the mixture was placed over whole mount retinas and incubated overnight at +4°C. Tissues were washed three times, 5 minutes each, with PBS after the overnight incubation of either α-SMA antibodies and mounted with mounting medium containing Hoechst 33258.

### Imaging and Analysis of Capillary Constrictions

Capillary constrictions were analyzed on fluorescein-lectin labeled whole-mount retinas. Primary branches of the central retinal artery were defined as the first-order retinal microvessels.

Vascular orders equal to or higher than 2 at the superficial retinal vascular layer were evaluated. Analyzed vessels were branches of a randomly selected first-order retinal vessel, whose entire network could be imaged uninterruptedly. Images were acquired using the combined tile-scan and Z-stack (10 layers each) modes of the Leica TCS SP8 confocal laser scanning microscope (25X/0.95 water objective). As the pericyte soma position with reference to the vessel wall significantly changes with the imaging plane, pericyte-associated diameters were measured at the juxtanuclear positions perpendicular to the vessel Ωaxis. Rarely, when the difference between values measured from two sides of a pericyte nucleus was relatively high, the mean of the two values was used. These values were divided by the initial diameter of each vascular segment for an intrinsic baseline correction, and this value was taken as the “juxtanuclear diameter ratio.” To ensure a homogeneous dataset, a single pericyte with a clearly visualized soma was attributed per vascular branch. Pericytes on the microvascular wall were identified based on their “bump-on-a-log” morphology and surrounding lectin-labeled basement membrane.

### 3D Modeling of Capillaries Ex vivo

For 3D reconstructions of capillaries, eight mice were used, including naïve (n=3), NA-treated (n=3), and NA and blebbistatin-treated (n=2) retinas. NA and blebbistatin were administered intravitreally, as described above. For these experiments, before eyeball extraction, we added a step of transcardial perfusion with heparin in saline, followed by 10% gelatin from porcine skin in PBS, at 42°C (*53*), to better preserve the intraluminal structure of capillaries. Eyeball and retina extraction was performed as described above. We also placed 0.17-mm-thick standard coverslip pieces over the glass slide as a placeholder between the top coverslip and the glass slide underneath, to prevent compression of whole-mount retina tissues. For 3D modeling of capillaries, sections double-stained with FITC-conjugated lectin lectin and α-SMA antibody were imaged using a 63X oil immersion objective (NA:1.4), at a resolution of 512×512 pixels, covering areas of 50×50 µm², with a Z-stack of approximately 50 µm and 0.3 µm Z-steps. Each stack was positioned to capture at least 40 µm of a capillary branch including a pericyte soma. These two-channel images were initially downsampled to 400×400 pixels for computational efficiency, followed by the application of a 1-pixel Gaussian filter. The stack was then resliced for isotropic voxel spacing of ∼0.125 µm. Deconvolution was performed using the Richardson-Lucy algorithm with DeconvolutionLab2, utilizing a computed point spread function. α-SMA and lectin channels were processed separately through all steps (*54*). Volumes were manually segmented for capillary lumen and α-SMA signals indicating pericyte processes and pericyte somata using 3DSlicer software. 3D surface reconstructions of capillary lumen in.stl format were processed with the Vascular Modeling Toolkit (http://www.vmtk.org). Centerlines of each capillary model were computed using the vmtkcenterlines command, and cross-sectional areas perpendicular to this centerline were generated and calculated with the vmtkcenterlinesections command.

Additionally, the centerline geometry parameter torsion was computed using the vmtkcenterlinegeometry command. Visualization of these capillary models, cross-sectional areas, and centerline, superimposed on α-SMA-labeled branches and pericyte somata, was performed using ParaView software. Cross-sectional areas were categorized into soma and outside soma areas. For areas outside the soma, only those at two lengths from the endpoint of the soma were considered. Cross-sectional areas and torsion measurements are along a segment are summarized by calculating the median along the whole segment or a specific zone of interest, unless indicated otherwise in the results. For torsion measurements, absolute values of positive and negative torsion values (in arbitrary units) were used. Pericyte process volumes associated with capillary torsional bends were manually selected using the polygon selection tool of ParaView software, and the volumes of these selected surfaces were computed for further comparisons.

### Two photon microscopy of retina

Experiments included 3 male BALB/c mice (2-3 months of age, ∼ 35g in body weight). Animals were housed in 12-hour dark/12-hour light cyclic conditions and fed ad libitum. All experiments were performed under general anesthetic using intraperitoneal injection of a ketamine (100mg/Kg) and xylazine (10mg/Kg) cocktail. Intraperitoneal injection of diluted fluorescein 0.1mL (1:4) was performed to allow visualization of retinal blood vessels.

In-vivo TPLSM retinal imaging was performed as previously described (*5, 55*). Anesthetized mice were stabilized with a bite bar on a custom-made set-up, allowing *in vivo* retinal imaging. Mouse body temperature was monitored during imaging (37 °C). The eyelids were removed, and a 6-0 suture was placed in the superior conjunctiva to gently rotate the eye. Using fine-tipped tweezers, the conjunctiva over the imaging area was removed, exposing the underlying sclera. A 5-mm-diameter coverslip was placed over the sclera, creating a planar surface necessary for retinal imaging below the sclera (field of view 400 × 400 μm) using a multiphoton microscope (Olympus FVMPERS; water objective 25x, NA=1.05) controlled by FV30S-SW software. For the excitation of fluorescein, we used a mode-locked Ti:sapphire (InSight; Spectra-Physics) set to 920nm.

Fifth-order capillaries or higher were identified by analyzing vessel branches through the entire volume of the retinal vascular plexuses. A longitudinal line scan in the central part of the capillary (zoom x3) was recorded (20000 cycles; 1.1 ms per cycle). This produces a distance/time image with a height of 20000 pixels, denoting the time period recorded and a width equal to the length of the longitudinal probe, indicating the capillary extent measured (0.331µm per pixel). As the red blood cells (RBCs) do not take up fluorescein, shadows surrounded by fluorescent signal in these recordings represent individual RBCs (Fig. 6B). The angle of the oblique shadow with respect to the horizontal axis allowed RBC speed to be calculated as speed=Δx/Δt (Fig.6C). For the capillary torsion component, volume stacks (z-stacks) were obtained before each blood flow measurement. A total of five measures for each component (volume & speed) were taken for each capillary.

Using ImageJ software, unbiased stereology was performed with home-made macros. A hundred dissectors (100-pixel height) were placed along the full recorded time period via systematic random sampling. The oblique shadow angles of the RBC were measured using the line tool in at least two locations within each dissector. Using an R-software script, we calculated the RBC speed through time. Capillary hemodynamics was determined without knowledge of capillary torsion calculations as they were performed by different users.

### Förster Resonance Energy Transfer (FRET)

FRET was employed in this study to demonstrate the close interaction between actin and the head of myosin heavy chain. In FRET experiments, the labeling order used in the double immunolabeling of α-SMA and Myosin11 was reversed, with the acceptor fluorophore-carrying antibody applied before the donor fluorophore-carrying antibody (*56*). Alexa Fluor-488 (donor) and Cy3 (acceptor) were selected as the FRET pair (*57*). Alexa Fluor-488-conjugated anti-rabbit IgG nanosecondary (Chromotek, srb488) (1:500 dilution in blocking solution, 1:350) was employed to label the primary antibody against Myosin11, combined with Cy3-conjugated antibody against α-SMA for enhancing resolution and interaction distance. An excitation-emission (ExEm) spectral unmixing approach was implemented to mitigate cross-excitation and cross-emission (spectral bleed through) artifacts encountered in intensity-based approaches (*30*). Donor-only, acceptor-only, and donor-acceptor samples were prepared simultaneously under the same conditions, without the use of the nuclear marker Hoechst 33258. All samples were imaged in the lambda-scan mode of Leica LasX software, using 488 nm and 552 nm lasers separately, with 5 nm wide windows of 3 nm steps within the range of 492-680 nm. The RT 15/85 beamsplitter for reflectance microscopy was used to avoid spectral losses filtered out by dichroic mirrors. Acquired spectral images were processed with Fiji and analyzed with the FRETTY plugin. Efficiency values are reported after subtracting the background value from the ROI value. High FRET efficiency was defined as above 0.8, in line with prior work on different biological models (*58–60*)

## Statistical Analysis

Descriptive statistics and tests were selected based on the evaluation of parametric assumption criteria. Hypothesis testing was conducted one-sided with a confidence interval of 95%. All statistical analyses were performed using IBM SPSS© 23 software.

For the analysis of the RNAscope dataset, signal data were grouped according to vascular order. Comparisons among all vascular orders were conducted using Kruskal–Wallis one-way analysis of variance, and for paired comparisons, the Mann-Whitney U test was utilized. The trend in transcript signals along the vascular branching orders was evaluated by the Jonckheere-Terpstra test. Correlation of transcript signals for the two probes was assessed using the Spearman correlation test.

In the analysis of the blebbistatin dataset, extreme outliers of JNDR, lying more than 3.0 times the interquartile range (IQR) below the first quartile or above the third quartile, were identified per vascular order group and removed from the initial dataset (5 out of 2116 values). The dataset was split up by vascular order, and Mann-Whitney U or Independent samples t-tests were employed depending on the result of parametric assumption testing, and one-tailed *p* values were presented.

## Acknowledgements

We would like to thank Jens Peter Gabriel for confocal image acquisition expertise in LasX Navigator, and Mesut Fırat for research animal care and maintenance. We also thank the Biological Optical Microscopy Platform (BOMP) of the University of Melbourne, Dr. Schienstock, and Prof. Mueller for assistance with two-photon microscopy.

## Author roles

GK performed *ex vivo* experiments and analysis, contributed to project planning, manuscript writing and figure preparation. NB performed, supervised, and contributed to project development regarding ISH experiments. MDB performed ISH experiments and analysis, and contributed to figure preparation. MH performed two-photon experiments and analysis, and contributed to figure preparation. LAM and SEE performed experiments and analysis regarding the generation of torsional forces, and contributed to figure preparation, and significantly contributed to manuscript writing and supervision. TD conceived the idea and designed the study; SEE and TD concepted and supervised the project, and manuscript writing.

## SUPPLEMENTARY DATA

**Supplementary Fig 1.**
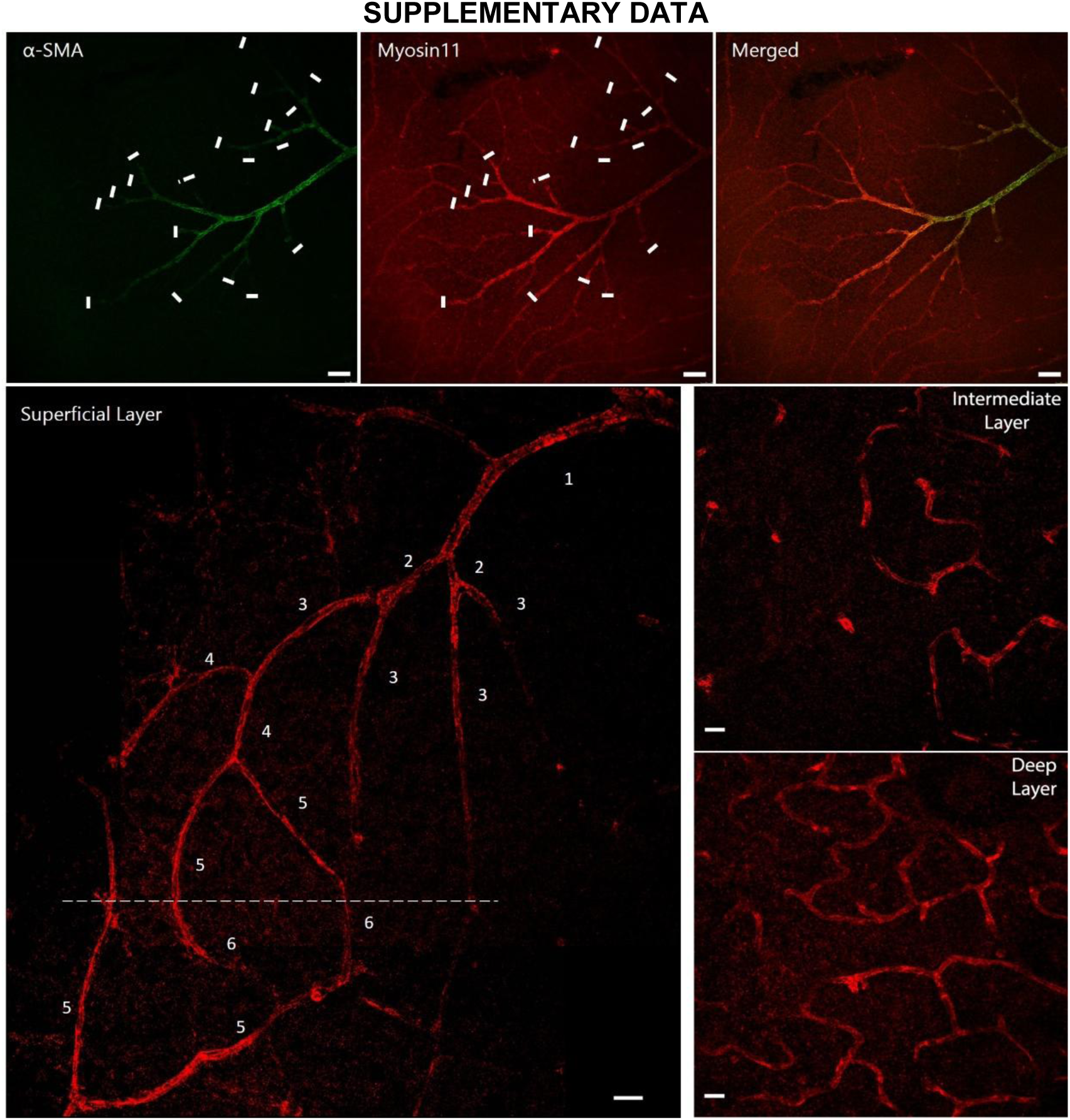
Unlike α-SMA, Myosin11 immunoreactivity is present throughout the retinal microvasculature. Upper row: Despite a significant reduction in α-SMA immunolabeling in downstream capillaries (green), Myosin11 immunostaining covers the entire microvasculature (red) in the superficial retinal vascular layer. The merged image highlights the disparity between the two immunolabeling. Of note, since a large area of retina was scanned using LasX Navigator, the dwell time per pixel was very short, which reduced the likelihood of detecting low α-SMA signals in downstream pericytes. Scalebar: 50 um. Lower row: Myh11 labeling persists along the entire superficial layer, revealing the capillary loops. The dashed line indicates the point where the capillary branches descend deeper to the intermediate vascular layer. The wide-field image was created by manually stitching together two Z-stack maximum projection images. Scalebar: 20 µm. Myh11 labeling uninterruptedly continues in the intermediate and deep layers of retinal vasculature. Scalebar: 20 µm. Z-stack maximum projection images of 8.88 and 8.45 µm thick vascular sections, respectively.

**Supplementary Figure 2:**
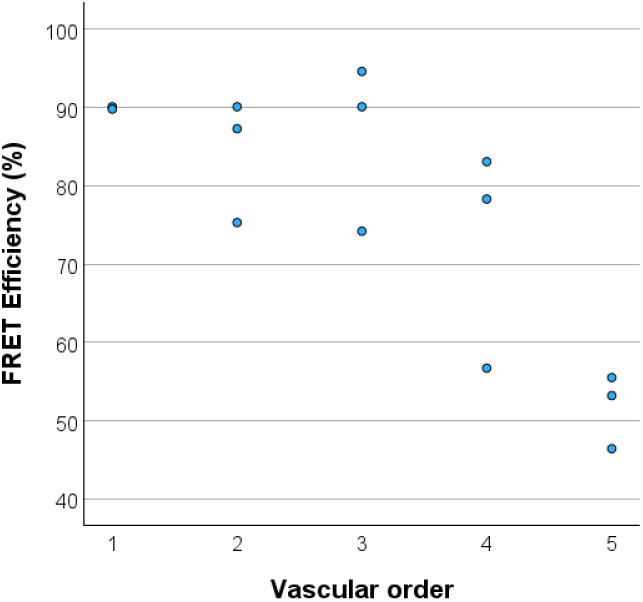
The plot presents the FRET efficiency averages over three ROIs (the green outlined regions in Figure 2C) in two or three retinal pericytes for each vascular order, as indicated (dots). Each dot represents the average value for each retinal pericyte evaluated.

**Supplementary Figure 3:**
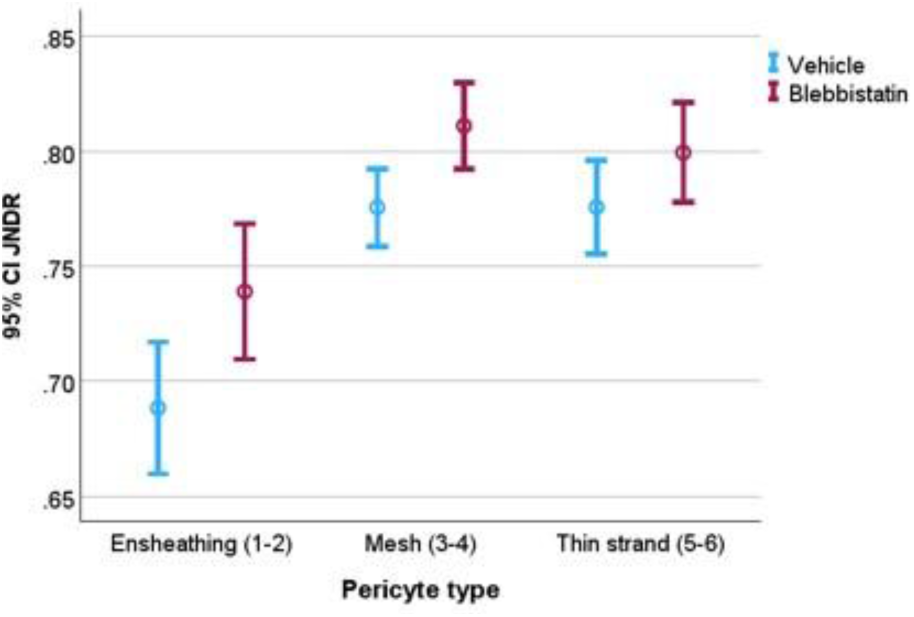
Graph illustrates that upstream and downstream pericyte contraction is suppressed by blebbistatin inhibition of actomyosin cross-bridge cycling. Due to high variation in vessel size across branching orders, for comparative analysis, the magnitude of constriction was normalized by calculating the ratio of the juxtanuclear radius to the radius at the origin of the capillary (referred to as juxtanuclear diameter ratio, JNDR). Blebbistatin or its vehicle was injected 40 minutes before NA. Both agents were administered intravitreally. Error bars represent 95% Confidence Interval (CI) for JNDR.

---

**Supplementary Table 1:**
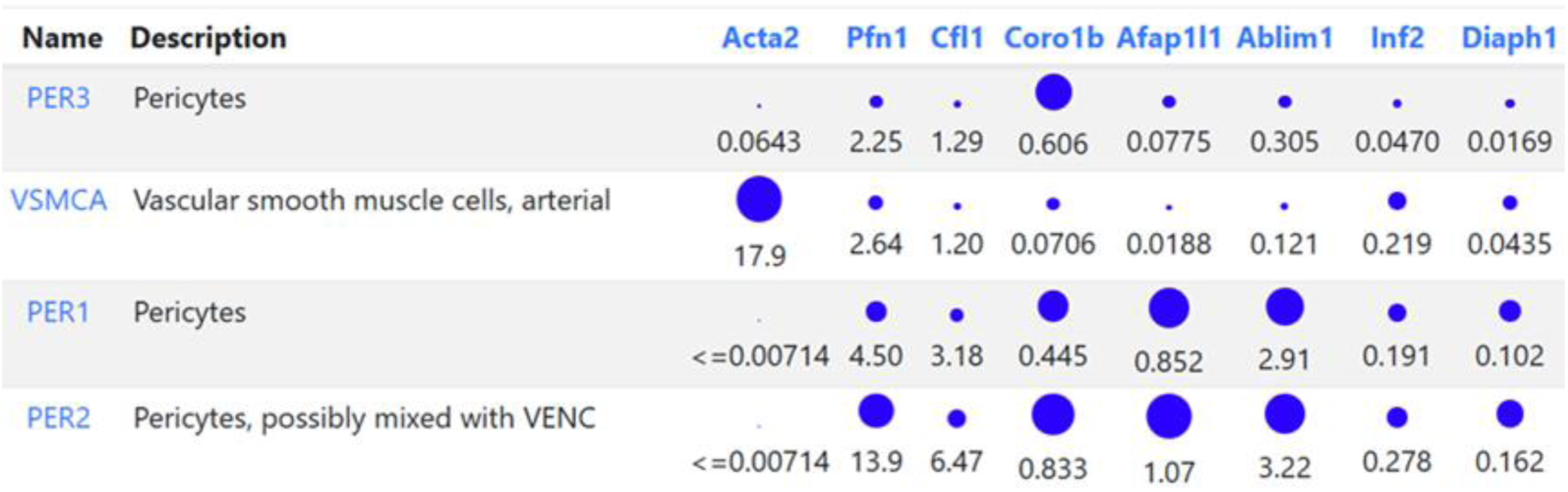
The importance of actin polymerization/depolymerization processes in pericytes is highlighted by the relative abundance of examples of related actin binding proteins compared to Acta2 (α-Smooth Muscle Actin gene). Adapted from mousebrain.org. Acta2: actin alpha 2, smooth muscle Pfn1: Profilin-1 Cfl1: Cofilin 1 Coro1b: Coronin 1B Afab1l1: actin filament associated protein 1 like 1 Ablim1: actin-binding LIM protein 1 Inf2: Inverted formin-2 Diaph1: diaphanous related formin 1.

